# Detection of early stage pancreatic cancer using 5-hydroxymethylcytosine signatures in circulating cell free DNA

**DOI:** 10.1101/422675

**Authors:** Francois Collin, Yuhong Ning, Tierney Phillips, Erin McCarthy, Aaron Scott, Chris Ellison, Chin-Jen Ku, Gulfem D Guler, Kim Chau, Alan Ashworth, Stephen R Quake, Samuel Levy

## Abstract

Pancreatic cancers are typically diagnosed at late stage where disease prognosis is poor as exemplified by a 5-year survival rate of 8.2%. Earlier diagnosis would be beneficial by enabling surgical resection or earlier application of therapeutic regimens. We investigated the detection of pancreatic ductal adenocarcinoma (PDAC) in a non-invasive manner by interrogating changes in 5-hydroxymethylation cytosine status (5hmC) of circulating cell free DNA in the plasma of a PDAC cohort (n=51) in comparison with a non-cancer cohort (n=41). We found that 5hmC sites are enriched in a disease and stage specific manner in exons, 3’UTRs and transcription termination sites. Our data show that 5hmC density is reduced in promoters and histone H3K4me3-associated sites with progressive disease suggesting increased transcriptional activity. 5hmC density is differentially represented in thousands of genes, and a stringently filtered set of the most significant genes points to biology related to pancreas (GATA4, GATA6, PROX1, ONECUT1) and/or cancer development (YAP1, TEAD1, PROX1, ONECUT1, ONECUT2, IGF1 and IGF2). Regularized regression models were built using 5hmC densities in statistically filtered genes or a comprehensive set of highly variable 5hmC counts in genes and performed with an AUC = 0.94-0.96 on training data. We were able to test the ability to classify PDAC and non-cancer samples with the Elastic net and Lasso models on two external pancreatic cancer 5hmC data sets and found validation performance to be AUC = 0.74-0.97. The findings suggest that 5hmC changes enable classification of PDAC patients with high fidelity and are worthy of further investigation on larger cohorts of patient samples.

## Introduction

Translational research using genomic and proteomic technologies has provided molecular insights into the pathogenesis and biology of pancreatic cancer but has yet to yield robust diagnostic biomarkers to impact early diagnosis of disease, as reflected by a low overall 5-year survival rate of 8.2%^1,2^. Pancreatic cancer often presents late and has few symptoms, at which point only 10-20% of patients are eligible for surgical resection ^2^. Pancreatic ductal adenocarcinoma (PDAC) and its variants constitute more than 90% of all pancreatic cancer cases ^3^ with the next most common sub-type being neuroendocrine tumors ^2^. Tobacco smoking confers a two- to three-fold higher risk of pancreatic cancer and also demonstrates a dose-risk relationship, while contributing to approximately 15 to 30% of cases ^2^, with smokers diagnosed 8 to 15 years younger than non-smokers^4,5^. Family history is contributory in approximately 10% of cases, and germline mutations in genes such as BRCA2, BRCA1, CDKN2A, ATM, STK11, PRSS1, MLH1 and PALB2 are associated with pancreatic cancer with variable penetrance ^2^.

The management of PDAC presents physicians with challenges along the entire clinical spectrum, including early detection in high risk individuals, early diagnosis of patients with symptoms or imaging findings, prognostication of outcomes and prediction of therapeutic responsiveness. Collectively these factors have engendered intensive efforts in translational research to identify and validate biomarkers with sufficient clinical performance metrics to improve decision algorithms and resultant clinical outcomes. Current guidelines in PDAC management are limited to two biomarker recommendations tested in an invasive fashion in cystic fluid. First, carbohydrate antigen 19-9 (CA 19-9) guides surgery decisions, use of adjuvant therapy, or the detection of post-operative tumor recurrence, however, the utility is limited because 10% of the patients does not secrete the antigen^6^. Second, carcinoembryonic antigen (CEA) concentration determination from cyst fluid is used to distinguish higher risk mucinous from non-mucinous cysts^7,8^, thereby mitigating risk. Among the inherited risk factors are genomic mutations such as BRCA2, which confers a 3.5-fold risk in carriers, with the probability of a germline mutation between 6 to 12% in PDAC patients with a first-degree relative diagnosed with PDAC^9^.

Molecular analyses of pancreatic cancer genomes have revealed activating mutations in KRAS and inactivation of CDKN2A, TP53 and SMAD4, either through point mutation or copy number changes at >50% population frequency^10–12^. However, mutational heterogeneity along with lack of disease specificity due to pleiotropy render this subset of genes incomplete for the diagnosis of patients. Molecular subtyping of pancreatic tumors using mutational-based data^11^ or gene expression signatures^13–15^ have not yet seen clinical applicability. Other forms of molecular profiling have focused on epigenetics, namely chromatin-based post-translation modifications and the methylation status of cytosine bases in DNA.

Control of DNA state and chromatin regulation have been observed to underpin the onset and progression of oncologic disease^16,17^. DNA methylation status of cytosine bases has been shown to associate with transcriptional regulation of gene expression. DNA methylation in promoters tends to associate with gene silencing whereas demethylation is associated with gene activation^18^. More recently, detailed understanding of demethylation has been enabled with precision around intermediate states during demethylation activation^19,20^. Specifically, discovery of TET enzyme-mediated methyl-cytosine oxidation to 5-hydroxymethyl cytosine (5hmC) has yielded novel signatures that have enabled the definition of cellular states ^21^, as well as the identification of cancer in the cell free state^22–24^.

Molecular signatures have been previously shown in circulating cell free DNA (cfDNA) based on 5-hydroxymethylation that may define the tissue of tumor origin in a variety of disease types^22^. Therefore, we embarked on a case-control study aimed at investigating whether DNA 5hmC signatures were present in the blood of PDAC patients compared to a cohort of non-cancer individuals. We also investigated whether these signatures enabled discrimination between cancer and non-cancer patients.

We found that in our study population, PDAC patients possess thousands of genes with altered hydroxymethylome compared to non-cancer individuals. Furthermore, filtering to those genes with the most differentially hydroxymethylated states, reveals genes that have been previously implicated in pancreas development or pancreatic cancer. This biologically significant gene set performs well in the construction of predictive models to discriminate PDAC from non-disease, suggesting that the measurement of 5hmC in cfDNA merits further investigation for the detection and classification of PDAC.

## Results

### Clinical cohort and Study Design

Plasma specimens from 92 subjects with or without pancreatic ductal adenocarcinoma (PDAC) were collected at multiple institutions in different geographic regions of the United States and Germany. These PDAC and non-cancer patient samples satisfied the study inclusion criteria, which included a minimum subject age of 18 years as well as confirmed pathologic diagnosis of adenocarcinoma of any subtype at the time of biopsy or surgical resection for subjects in the cancer cohort (Figure 1A). The non-cancer cohort was identified as satisfying the study inclusion criteria and patients were specifically negative for any form of cancer. Neither cohort were being treated with medication for disease at the time of blood collection, which was prior to any biopsy or surgical resection in the cancer cohort. There were no statistically significant differences in subject age or gender between the two cohorts, but there was a statistically significant greater tobacco exposure in the PDAC cohort, as expected given smoking is a common risk factor in pancreatic cancer.

**Figure 1:**
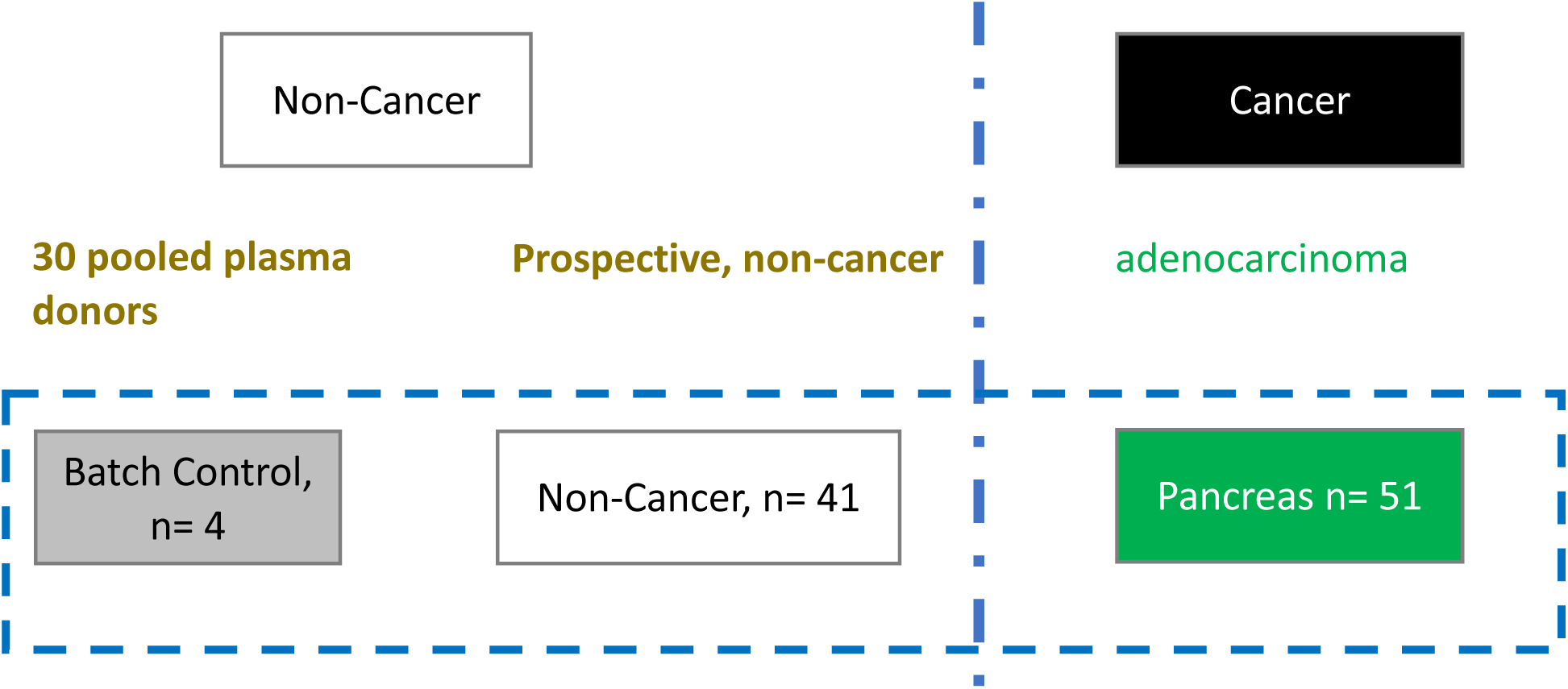
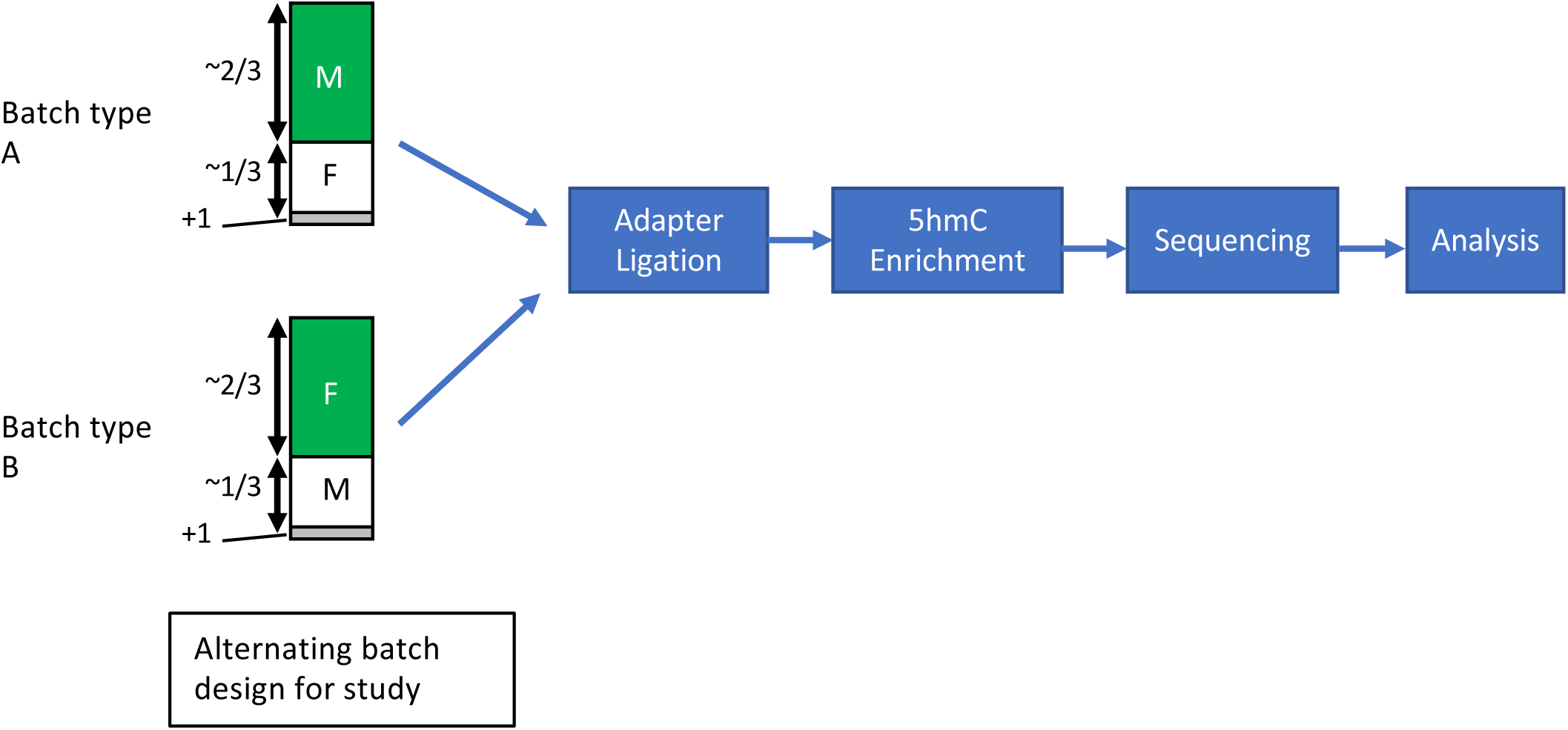
Study Design and Patient Cohorts Employed. A. Schematic depicting study cohorts employed. PDAC, n = 51, Non-cancer, n=41 and pooled non-cancer replicates were included across multiple 5hmC assay processing and sequencing batches. B. Schematic depicting sample processing workflow incorporating alternating flowcell constructs according to subject sex for detection of sample swaps.

### Sequencing results and metrics

PDAC samples that were exclusively either male or female were combined with non-cancer samples that were exclusively either female or male in a ratio of 2:1 respectively following a block randomization scheme. This generated two batch types for the study (Figure 1B) and enabled the identification of sample swaps, none of which were detected. A single pooled non-cancer sample (30 non-cancer donors were combined into one plasma pool from which cfDNA was isolated) served as a technical/process control for the each of the batches in the study. 5hmC enrichment libraries were sequenced to produce a median number of unique read pairs of 9.1 and 10.7 million in the PDAC and non-cancer cohorts respectively. Filtering criteria to enable the determination of high quality 5hmC libraries were established yielding 51 PDAC and 41 non-cancer subjects. These criteria were established from previous studies^22^ and extensive analysis did not reveal batch processing effects occurring specifically in either study cohort.

### Distributions of 5hmC densities into functional regions in PDAC and non-cancer cohort

To gain an understanding of the genomic regions associated with hydroxymethylation, we first determined 5hmC enriched loci, as measured by increased read density and detection as peaks by MACS2. The vast majority of 5hmC loci occur on average in non-coding regions of the genome (intronic, transposon repeats – SINES and LINEs, and intergenic - Figure 2A) with no preferential distribution in the PDAC or non-cancer cohort. Despite the high frequency of 5hmC sites, these functional regions exhibit low enrichment (intron, 2B) or even depletion of 5hmC (intergenic and LINE elements, 2B) relative to the genome background. Instead, enrichment occurs in promoters, UTRs, exons, transcription termination sites (TTS) and SINE elements, as measured by increased relative fold change compared to the genome background. Significant differences in enrichment of 5hmC peaks over functional regions were observed in a disease cohort specific manner. Increases in 5hmC peak enrichment in PDAC was measured in exons, 3’UTR and TTS whereas decreases were found in promoter and LINEs, which themselves were either enriched or depleted respectively (Figure 2C). These global changes, found to occur in a statistically significant manner in each cohort, were also detected in a cancer stage specific manner, with gradual increases (exon, 3’UTR and TTS) or decreases (promoter and LINE) in later stage patients (Figure 2D).

Next, we investigated 5hmC occupancy, and its associated changes in PDAC, with respect to chromatin state. Post-translational modifications such as methylation and acetylation on histone proteins were inferred in relation to 5hmC occupancy, using the existing histone maps from the pancreatic cancer cell line, PANC-1, for which epigenetic data were made available by ENCODE^25^. Notably, reduced overlap with 5hmC was observed in PDAC coincident with loci associated with H3K4me3 and H3K27ac, both of which mark transcriptionally active states (Figure 2E). There was no significant global difference between PDAC and non-cancer cohorts in 5hmC peak overlap with H3K4me1-associated loci, which mark enhancer regions (Figure 2E). Furthermore, 5hmC peak overlap with H3K4me3-associated loci is significantly reduced with progressive disease as observed when the PDAC cohort is subdivided into disease stages (p=0.0003 – Figure 2E bottom right panel). This would be consistent with increased expression of the actively transcribed genes in PANC-1 cancer cell line, in the PDAC cohort compared to non-cancer cohort. Conversely, there were no statistically significant changes detected globally in 5hmC peak occupancy over H3K4me1- and H3K27ac-associated loci with progressive disease (Figure 2E – bottom panels). Amongst the three histone maps from PANC-1 cell lines, H3K4me1 has the most similarity with 5hmC occupancy in both the PDAC and non-cancer cohort whereas H3K4me3 histone map has the lowest similarity, as calculated by Jaccard index (Supplementary Figure 1). Moreover, 5hmC read density is highest at the peak center for H3K4me1-associated sites in general whereas it is lowest at the peak center for H3K4me3-associated sites (Figure 2F). Overall, 5hmC exhibits diverse patterns over these different chromatin contexts, which may potentially be indicative of varied interplay between hydroxymethylation and different histone modifications.

**Figure 2.**
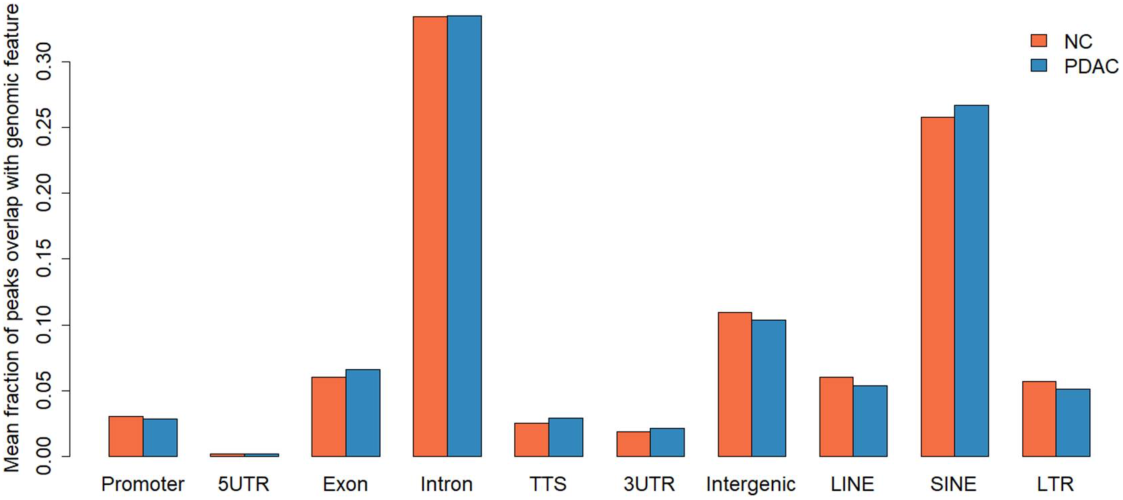
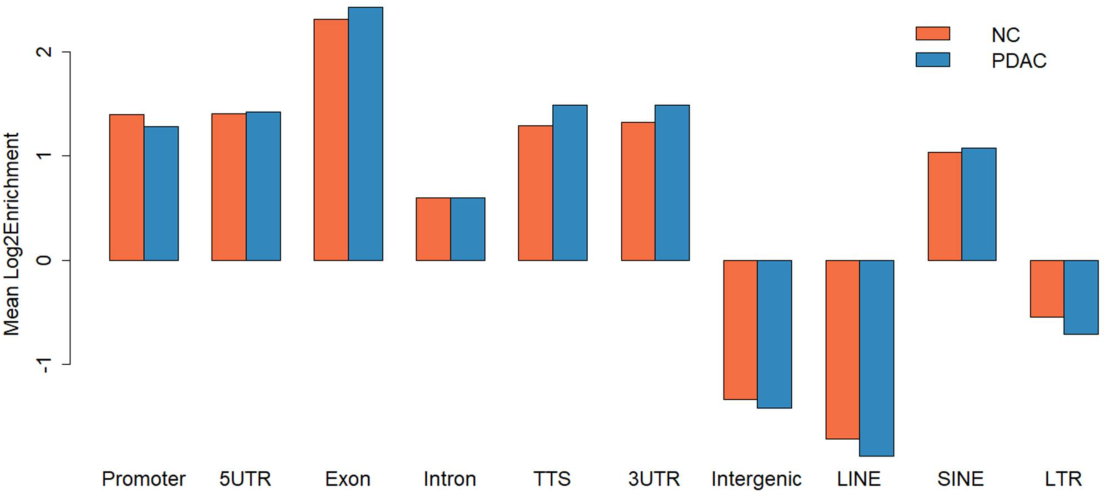
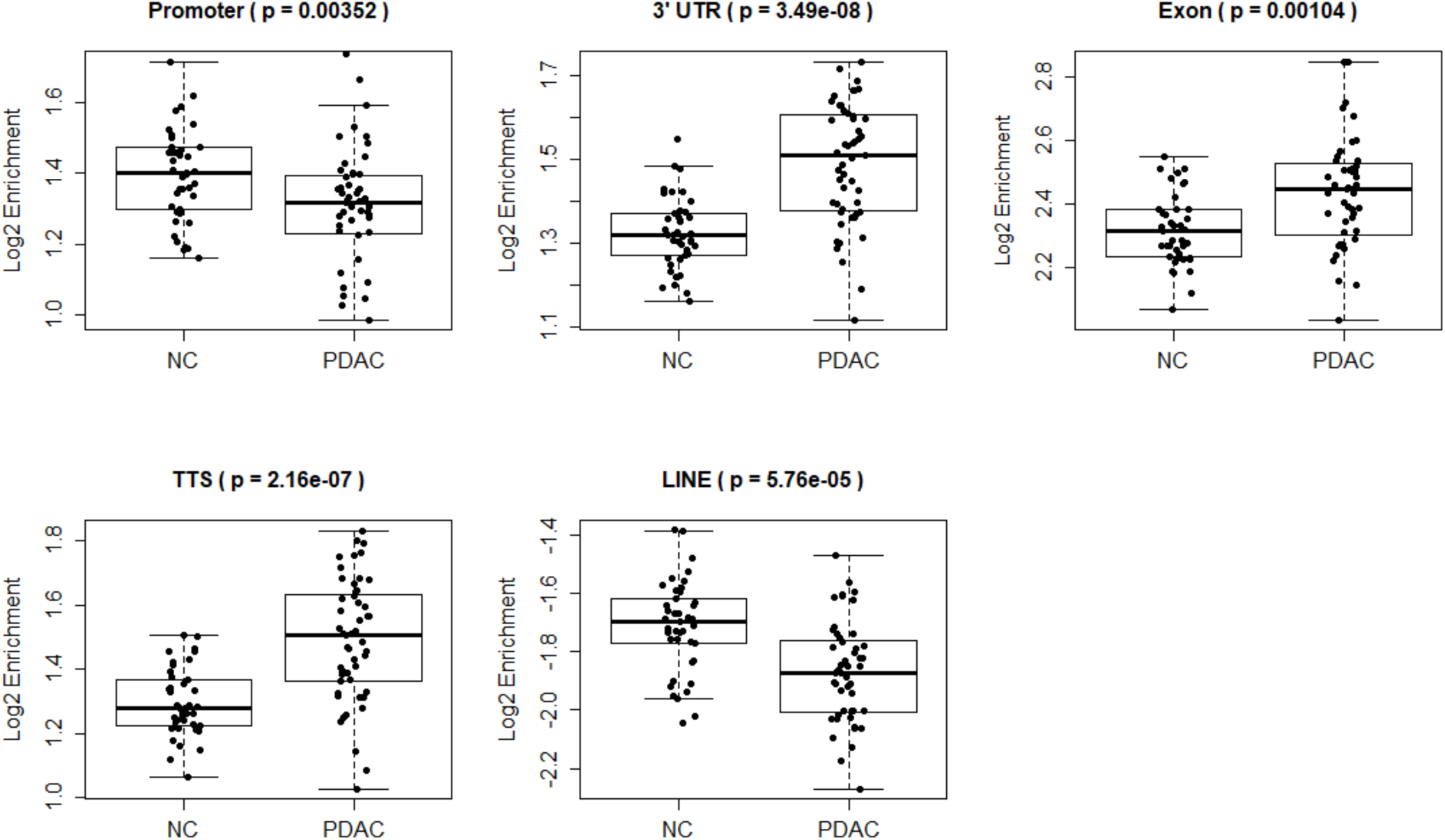
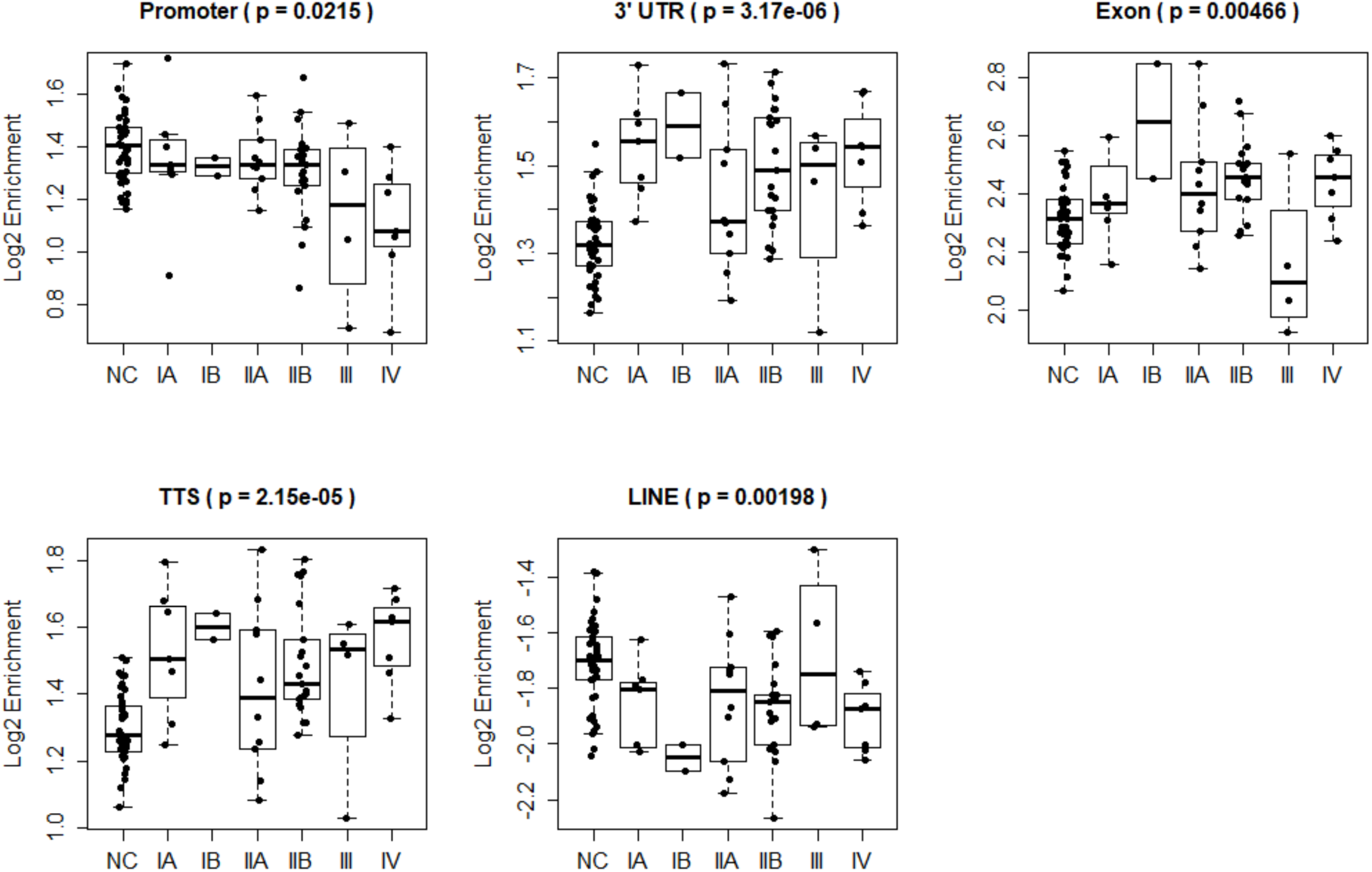
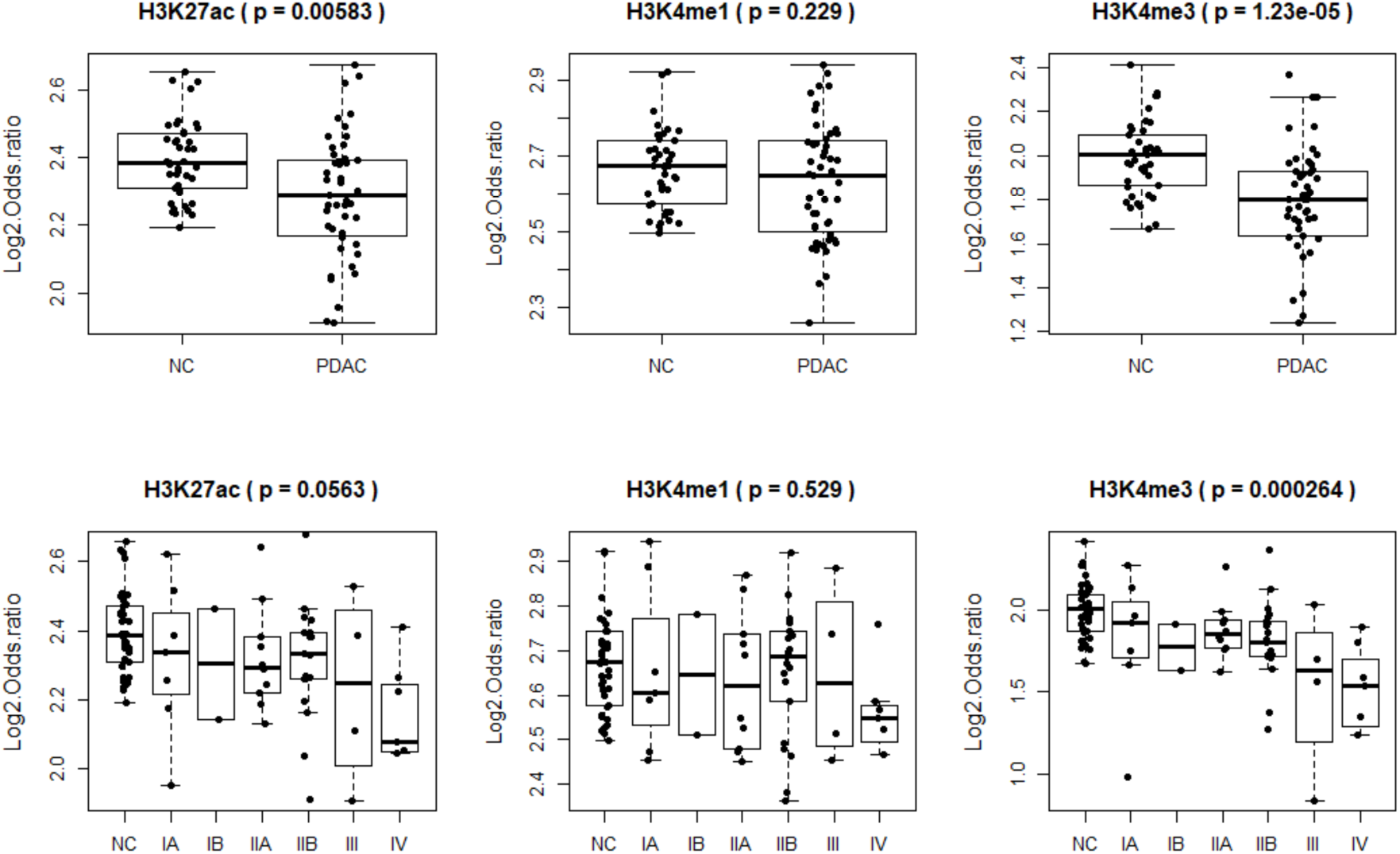
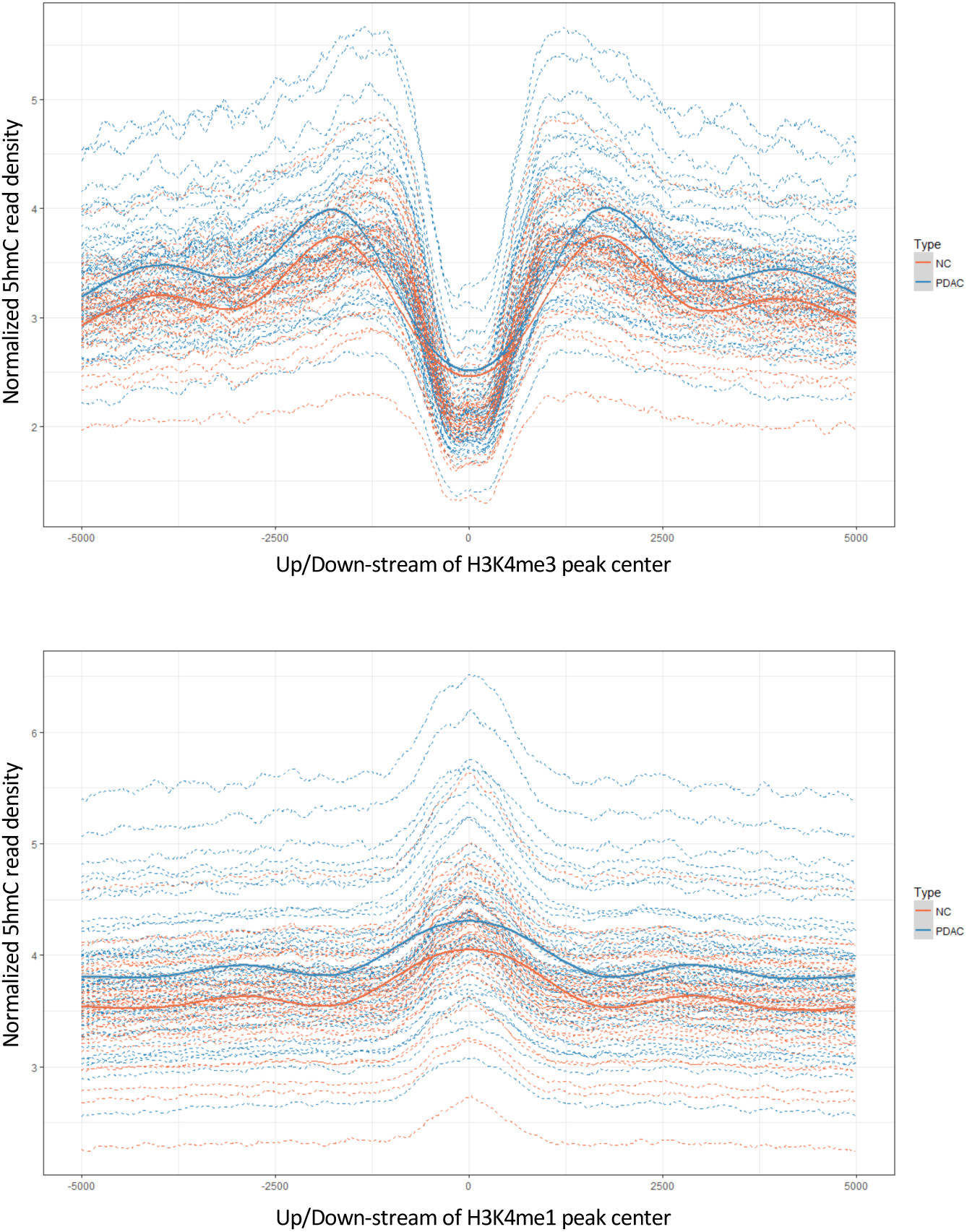
Differential enrichment of 5hmC in functional genomic regions in PDAC compared with non-cancer (NC) samples. A. 5hmC peak distribution in genomic features. Note non-coding features have larger number of peaks. B. Enrichment analysis (Y-axis = log2 (PDAC/non-cancer)) shows that gene-based features, SINEs and Alus are enriched in 5hmC peaks in both PDAC and non-cancer cohorts. Intergenic, LINEs and L1s are depleted of 5hmC peaks. C. Box plots depicting statistically significant changes of 5hmC peaks in promoter and LINE elements in pancreas cancer samples (decreased accumulation). Exons, 3’UTR and translation termination sites are enriched in cancer samples. Y-axis = log2 (PDAC/non-cancer) D. Box plots depicting statistically significant changes of 5hmC peaks in functional genomic regions across pancreatic cancer stages. E. Box plots depicting 5hmC occupancy in PDAC and non-cancer (NC) cohorts over H3K27ac, H3K4me1 and H3K4me3 histone sites from PANC-1 cell line data obtained from ENCODE. Note 5hmC peak depletion in H3K4me3 and H3K27ac histone marks in the PDAC cohort (top panel) and H3K4me3 depletion with progressive disease (bottom panel). F. Normalized 5hmC read density in the PDAC and non-cancer (NC) cohorts over 10kb windows surrounding H3K4me3- (top panel) and H3K4me1- (bottom panel) associated sites as determined in PANC-1 cell line. Each individual sample is depicted with a dotted line: Dotted red lines == Non-cancer (NC) sample, dotted blue lines = PDAC sample. Solid lines show average intensity of normalized 5hmC counts across all non-cancer(NC) samples (solid red line) or all PDAC samples (solid blue line).

### Identification of disease specific genes from plasma samples

Differential analysis of 5hmC densities in genes revealed 6,496 and 6,684 genes with an increased and decreased 5hmC density respectively in PDAC compared to non-cancer samples (Figure 3A – adjusted p-value < 0.05). Further filtering of this gene set (fold change ≥ 1.5 in PDAC versus non-cancer and average log 2 CPM ≥ 4 counts, 142 genes total) revealed annotated genes with increased 5hmC density and whose biology is related to pancreas development (GATA4^26^, GATA6^26^, PROX1^27^, ONECUT1^28^) and/or implicated in cancer (YAP1^29^, TEAD1^29^, PROX1^30^, ONECUT2/ONECUT1, IGF1 and IGF2). Inspection of the MSigDB for relevant pathways comprising the 142 genes with enriched 5hmC densities revealed a preponderance of pathways down-regulated in liver cancer (5 of the top 10 most significant pathways – Table 2). The differential representation analysis coupled with filtering (fold change ≤ 1.5 in PDAC versus non-cancer and log CPM of 5hmC ≥ 4) also revealed 178 genes with a decreased 5hmC density in PDAC cfDNA. Closer inspection of these pathways with decreased 5hmc representation revealed fundamental pathways in immune system regulation (3 of the top 10 most significant pathways – Table 3).

**Table 1.** Clinical Characteristics of Non-Cancer and Cancer Subject Cohorts.

|  | Non-Cancer | Cancer |
| --- | --- | --- |
| **Age<sup>+</sup>** |  |  |
|  | 66.0 | 71.2 |
| **Gender(%)** |  |  |
| Male | 60.0 | 45.1 |
| **Smoking History** |  |  |
| Status(%) |  |  |
| Current | 19.5 | 19.6 |
| Former | 29.3 | 37.3 |
| None | 51.2 | 43.1 |
| **Pack-Years<sup>+</sup>** |  |  |
| Current | 5.3 | 29.6 |
| Former | 20.5 | 24.2 |
| None | NA | NA |
| **Pack-Years<sup>+</sup>** |  |  |
| All | 14.4 | 25.7 |
| **Time Since Cessation<sup>+</sup>** |  |  |
| Month | 264.2 | 272.3 |
| **Stage(%)** |  |  |
| I | NA | 18 |
| II | NA | 61 |
| III | NA | 7.8 |
| IV | NA | 14 |

+ mean of Non-Cancer and Cancer groups.
Other values are percentages of each category in “Non-Cancer” and “Cancer” groups.

**Table 2.**
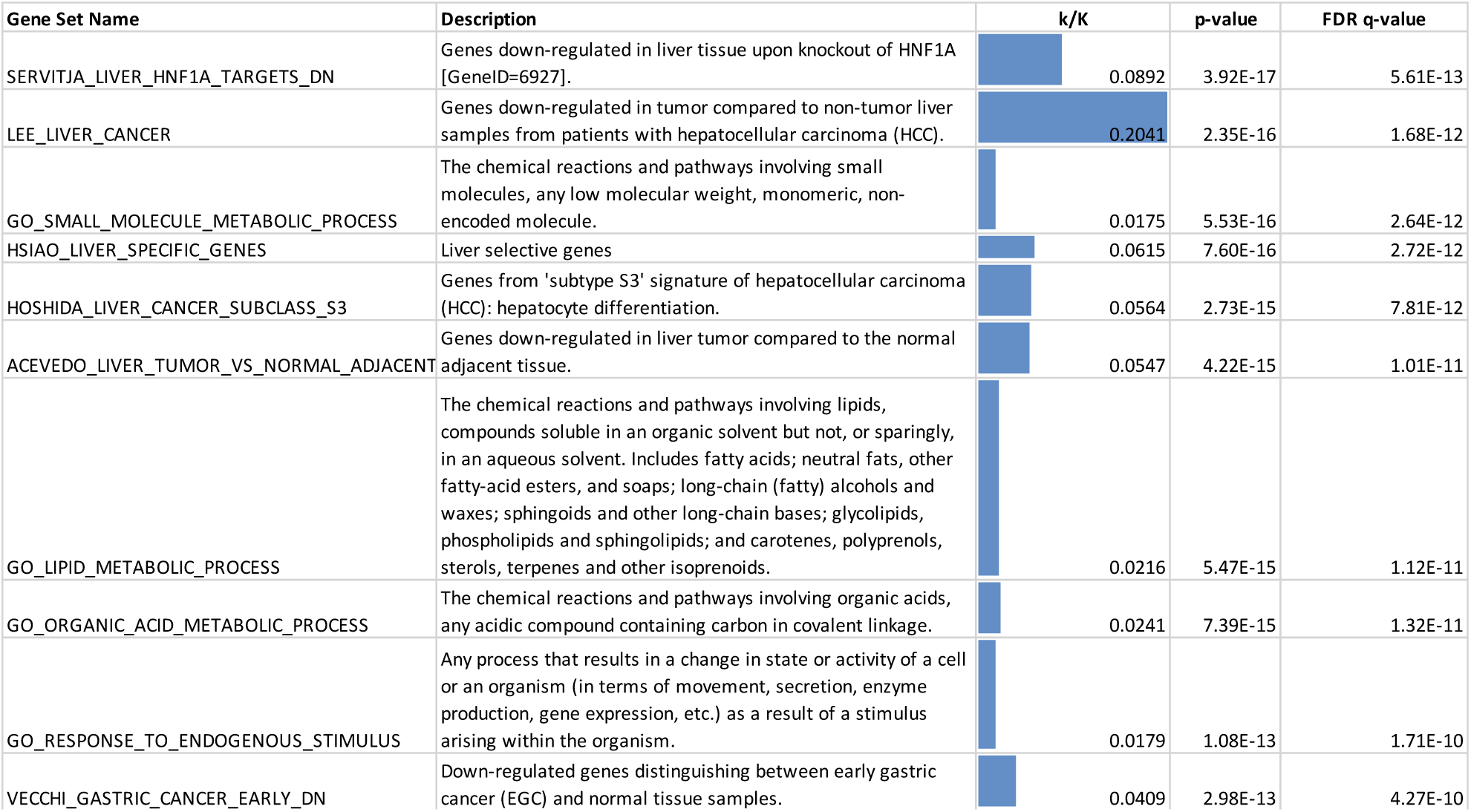
Top 10 pathways represented by 142 genes with increased 5hmC density in PDAC samples versus non-cancer samples.

| Gene Set Name | Description | k/K | p-value | FDR q-value |
| --- | --- | --- | --- | --- |
| SERVITJA_LIVER_HNF1A_TARGETS_DN | Genes down-regulated in liver tissue upon knockout of HNF1A [GeneID=6927]. | 0.0892 | 3.92E-17 | 5.61E-13 |
| LEE_LIVER_CANCER | Genes down-regulated in tumor compared to non-tumor liver samples from patients with hepatocellular carcinoma (HCC). | 0.2041 | 2.35E-16 | 1.68E-12 |
| GO_SMALL_MOLECULE_METABOLIC_PROCESS | The chemical reactions and pathways involving small molecules, any low molecular weight, monomeric, non-encoded molecule. | 0.0175 | 5.53E-16 | 2.64E-12 |
| HSIAO_LIVER_SPECIFIC_GENES | Liver selective genes | 0.0615 | 7.60E-16 | 2.72E-12 |
| HOSHIDA_LIVER_CANCER_SUBCLASS_S3 | Genes from 'subtype S3' signature of hepatocellular carcinoma (HCC): hepatocyte differentiation. | 0.0564 | 2.73E-15 | 7.81E-12 |
| ACEVEDO_LIVER_TUMOR_VS_NORMAL_ADJACENT | Genes down-regulated in liver tumor compared to the normal adjacent tissue. | 0.0547 | 4.22E-15 | 1.01E-11 |
| GO_LIPID_METABOLIC_PROCESS | The chemical reactions and pathways involving lipids, compounds soluble in an organic solvent but not, or sparingly, in an aqueous solvent. Includes fatty acids; neutral fats, other fatty-acid esters, and soaps; long-chain (fatty) alcohols and waxes; sphingoids and other long-chain bases; glycolipids, phospholipids and sphingolipids; and carotenes, polyprenols, sterols, terpenes and other isoprenoids. | 0.0216 | 5.47E-15 | 1.12E-11 |
| GO_ORGANIC_ACID_METABOLIC_PROCESS | The chemical reactions and pathways involving organic acids, any acidic compound containing carbon in covalent linkage. | 0.0241 | 7.39E-15 | 1.32E-11 |
| GO_RESPONSE_TO_ENDOGENOUS_STIMULUS | Any process that results in a change in state or activity of a cell or an organism (in terms of movement, secretion, enzyme production, gene expression, etc.) as a result of a stimulus arising within the organism. | 0.0179 | 1.08E-13 | 1.71E-10 |
| VECCHI_GASTRIC_CANCER_EARLY_DN | Down-regulated genes distinguishing between early gastric cancer (EGC) and normal tissue samples. | 0.0409 | 2.98E-13 | 4.27E-10 |

**Table 3.**
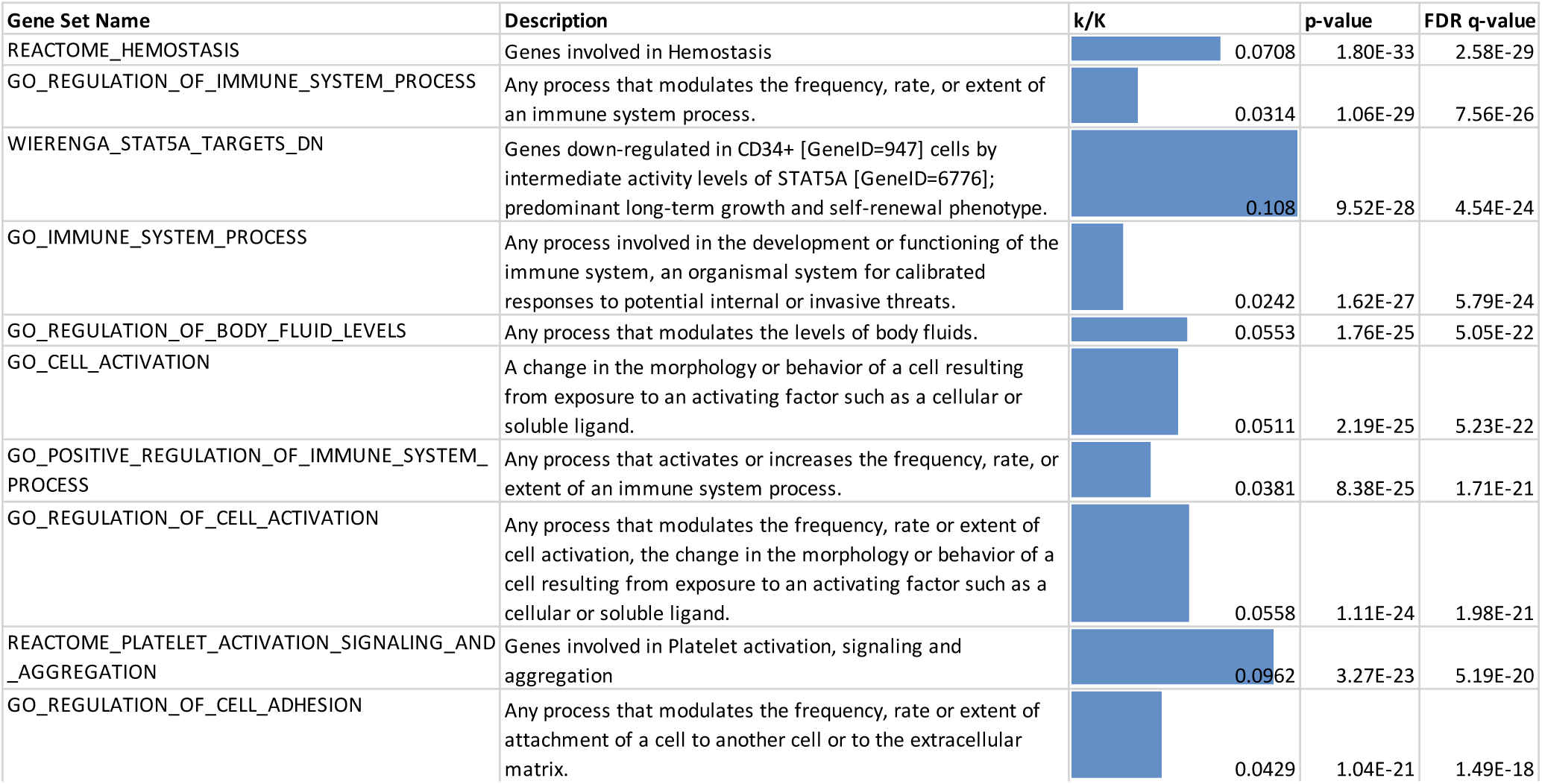
Top 10 pathways represented by 178 genes with decreased 5hmC density in PDAC samples versus non-cancer samples.

| Gene Set Name | Description | k/K | p-value | FDR q-value |
| --- | --- | --- | --- | --- |
| REACTOME_HEMOSTASIS | Genes involved in Hemostasis | 0.0708 | 1.80E-33 | 2.58E-29 |
| GO_REGULATION_OF_IMMUNE_SYSTEM_PROCESS | Any process that modulates the frequency, rate, or extent of an immune system process. | 0.0314 | 1.06E-29 | 7.56E-26 |
| WIERENGA_STAT5A_TARGETS_DN | Genes down-regulated in CD34+ [GeneID=947] cells by intermediate activity levels of STAT5A [GeneID=6776]; predominant long-term growth and self-renewal phenotype. | 0.108 | 9.52E-28 | 4.54E-24 |
| GO_IMMUNE_SYSTEM_PROCESS | Any process involved in the development or functioning of the immune system, an organismal system for calibrated responses to potential internal or invasive threats. | 0.0242 | 1.62E-27 | 5.79E-24 |
| GO_REGULATION_OF_BODY_FLUID_LEVELS | Any process that modulates the levels of body fluids. | 0.0553 | 1.76E-25 | 5.05E-22 |
| GO_CELL_ACTIVATION | A change in the morphology or behavior of a cell resulting from exposure to an activating factor such as a cellular or soluble ligand. | 0.0511 | 2.19E-25 | 5.23E-22 |
| GO_POSITIVE_REGULATION_OF_IMMUNE_SYSTEM_PROCESS | Any process that activates or increases the frequency, rate, or extent of an immune system process. | 0.0381 | 8.38E-25 | 1.71E-21 |
| GO_REGULATION_OF_CELL_ACTIVATION | Any process that modulates the frequency, rate or extent of cell activation, the change in the morphology or behavior of a cell resulting from exposure to an activating factor such as a cellular or soluble ligand. | 0.0558 | 1.11E-24 | 1.98E-21 |
| REACTOME_PLATELET_ACTIVATION_SIGNALING_AND_AGGREGATION | Genes involved in Platelet activation, signaling and aggregation | 0.0962 | 3.27E-23 | 5.19E-20 |
| GO_REGULATION_OF_CELL_ADHESION | Any process that modulates the frequency, rate or extent of attachment of a cell to another cell or to the extracellular matrix. | 0.0429 | 1.04E-21 | 1.49E-18 |

Expanding gene set enrichment analysis to include the full data set of all genes revealed that more than 30% of immune related pathways have a reduced 5-hydroxymethylation across early and late stage PDAC (Figure 3 B, Table 4). Multidimensional scaling analysis (MDS) using either the 13,180 genes with high variation in 5hmC counts (Figure 3 C) or the 320 genes filtered at the extremes of 5hmC representation in PDAC (Figure 3 D), reveal partitioning of the PDAC samples from the non-cancer equally well. Furthermore, hierarchical clustering using the 320 genes revealed better partitioning of the 5hmC data from PDAC and non-cancer cfDNA from Song et al^22^ compared with similar data from Li et al^24^ (Figure 3E). In summary, we have been able to find a differentially represented gene set whose biological functions are congruent with both pancreatic development and cancer more broadly. Furthermore, 5-hydroxymethylation densities of these genes alone enable the partitioning of PDAC from non-cancer.

**Table 4.** MSigDB pathways containing genes with modification in 5hmC in PDAC. Down = number of pathways with genes that have reduced 5hmC in PDAC, Up = number of pathways with genes that have increased 5hmC in PDAC, [Up, Down]/Total.pathway = Down and Up values expressed as a ratio. Hallmark =, C2 = Curated gene sets inclusive of Biocarta, KEGG and Reactome, C5.BP = GO Biological processes, C6 = Oncogenic signatures, C7 = Immunologic signatures. Note that largest magnitude of change in the most gene rich set is a decreased 5hmC in immunologic genes.

|  | Total.pathway* | Down | Up | Down/Total.pathway | Up/Total.pathway |
| --- | --- | --- | --- | --- | --- |
| Hallmark | 50 | 18 | 14 | 0.360 | 0.280 |
| C2 | 3762 | 566 | 167 | 0.150 | 0.044 |
| C5.BP | 3426 | 251 | 123 | 0.073 | 0.036 |
| C6 | 187 | 11 | 0 | 0.059 | 0.000 |
| C7 | 4782 | 1545 | 287 | 0.317 | 0.059 |

**Figure 3.**
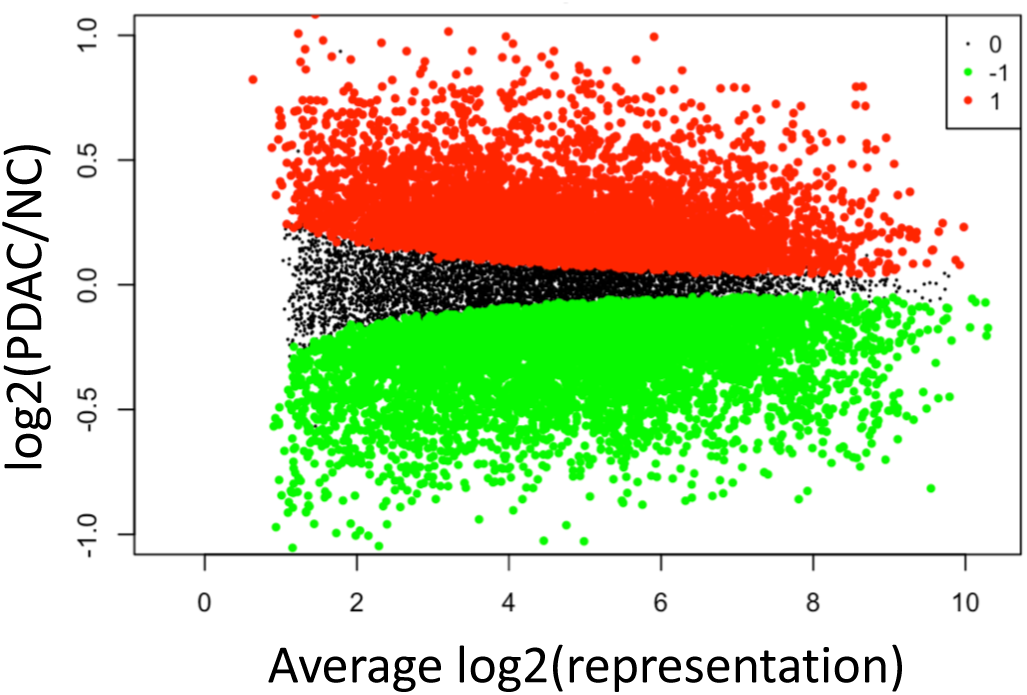
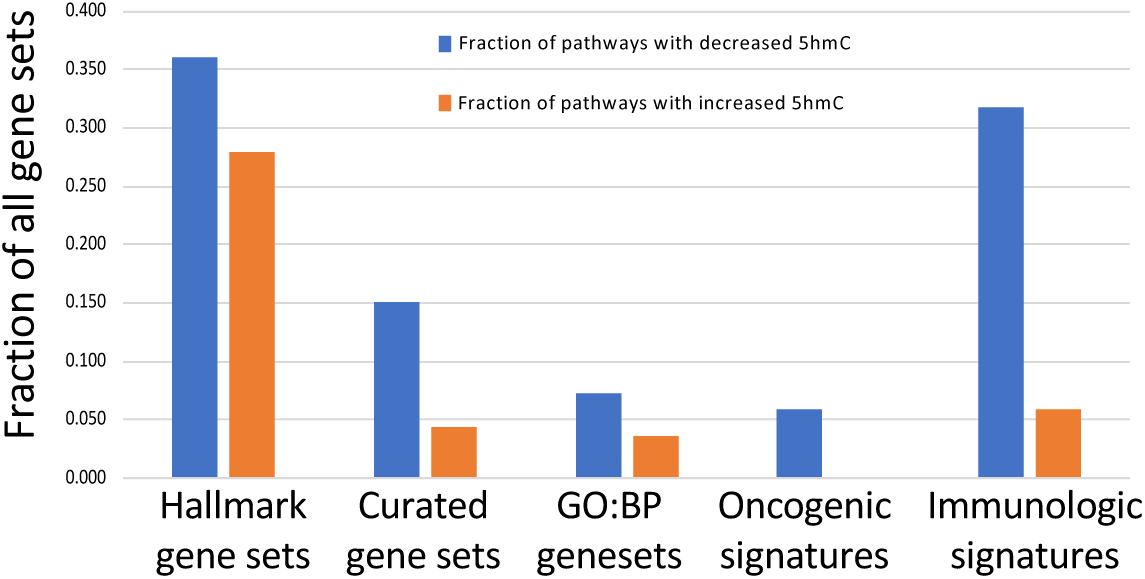
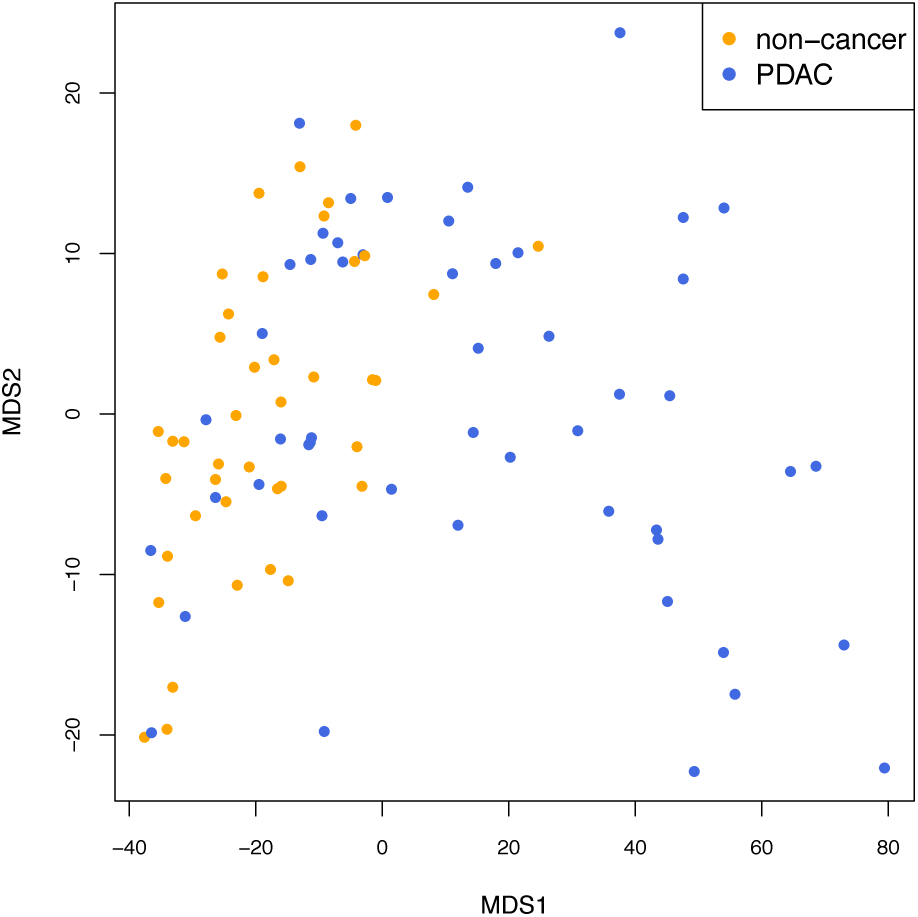
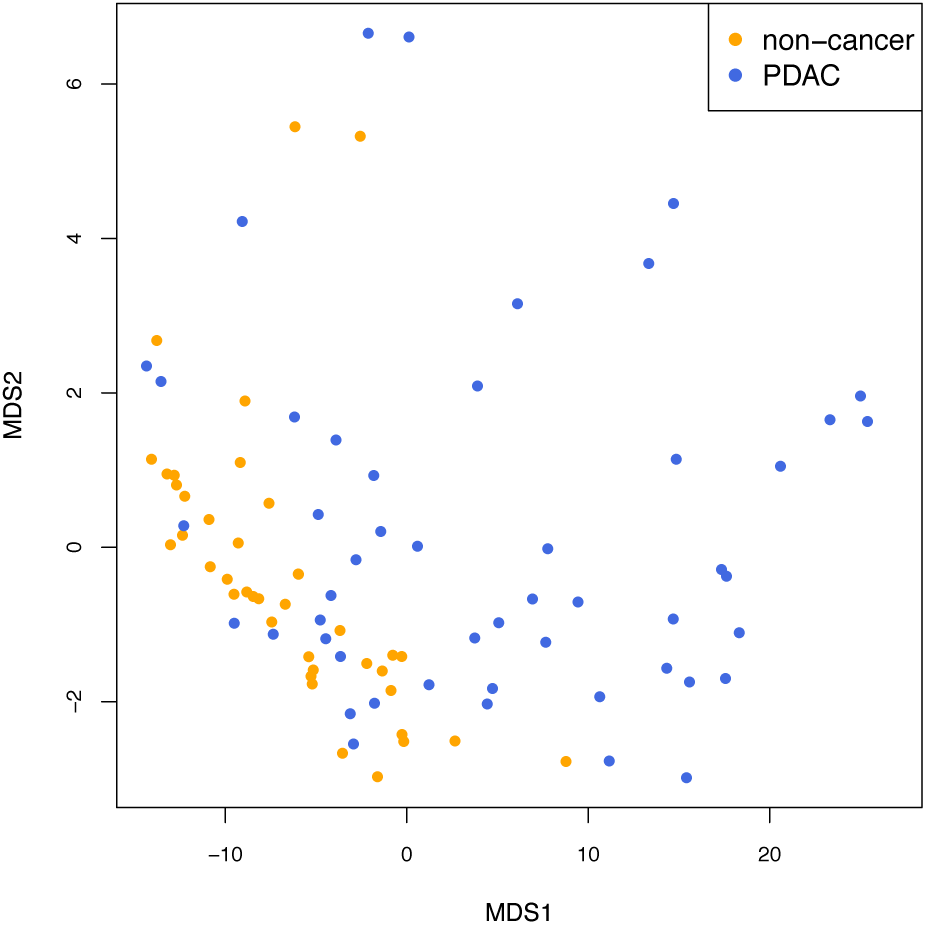
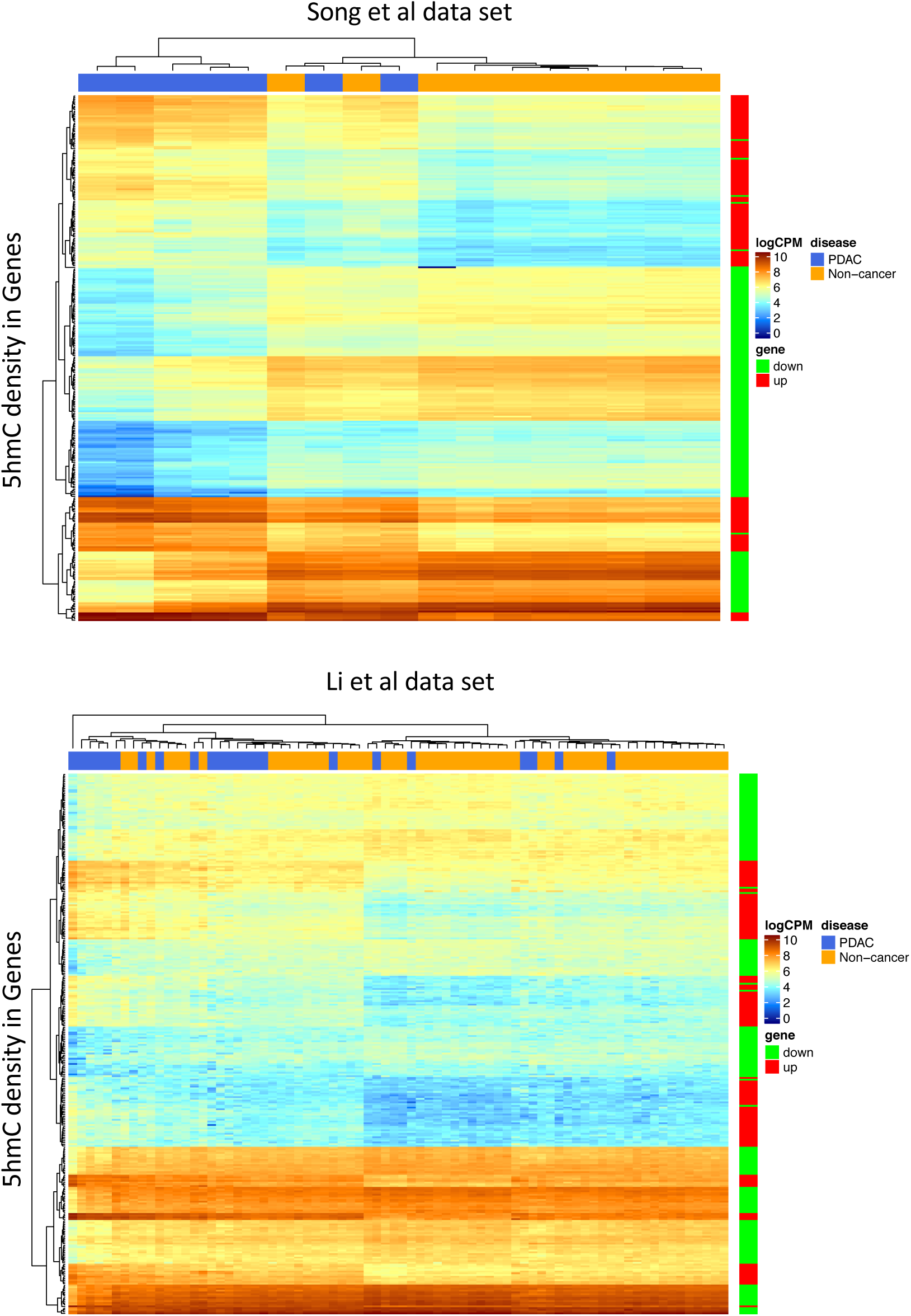
Identification of statistically significant 5hmC changes in genes in the PDAC cohort, biological significant of the gene set and ability to partition between PDAC and non-cancer samples using 5hmC counts. A. MA-Plot showing all differentially represented genes. Adjusted p-value < 0.05, NC = non-cancer cohort. Red points mark genes with increased 5hmC density in PDAC versus NC. Green points mark genes with decreased 5hmC density in PDAC versus NC. Red and Green are significant at the adjusted p-value = 0.05. B. GSEA using differentially 5hmC enriched genes reveals >20% KEGG pathways are both up and down-represented via 5hmC changes in pancreas cancer versus non-cancer samples. Also >30% immune pathways are down-represented in pancreas cancer versus non-cancer samples. C. MDS using log (counts per million) of 13,180 genes with statistically significant (adjusted p-value < 0.05) increase or decrease in 5hmC. Note reasonable partitioning of PDAC from non-cancer samples. D. PCA using log (counts per million) of 320 genes with statistically significant (adjusted p-value < 0.05) and filtered for increased PDAC representation (| log2(5hmC-PDAC/5hmC-non-cancer) | >= 0.58 and log2(average representation) >= 4) increase or decrease in 5hmC. Note reasonable partitioning of PDAC from non-cancer samples despite an order of magnitude smaller gene set than Figure 3 C. E. Heatmaps depicting hierarchical clustering results employing 320 genes (rows in heatmaps) to show how labelled samples (columns in heatmap) can be partitioned using log(CPM) 5hmC counts. Note almost perfect partitioning of the Song et al data set (top heatmap) versus incomplete partitioning of the Li et al data set (bottom heatmap).

### Predictive models for the detection of pancreatic cancer in cfDNA

We performed regularized logistic regression analysis in order to determine whether gene-based features are present in the PDAC and non-cancer cohorts that can enable the classification of patient samples. The full set of 92 patient samples were partitioned into a training and test set comprising 75% and 25% of the patient data respectively. We employed 65% of the genes with the most variable 5hmC density for model selection. Two methods of regularization, Elastic net (*glmnet*) and Lasso (*glmnet2*)^31^, were utilized. Other modeling approaches, such as random forest, support vector machines and neural networks, were explored in a preliminary analysis and were found to have inferior performance on the training data.

Both Elastic net (*glmnet*) and Lasso (*glmnet2*)^31^ regularization methods require specifying hyper-parameters which control the level of regularization used in the fit. These hyper-parameters were selected based on out-of-fold performance on 30 repetitions of 10-fold cross-validated analysis of the training data. Out-of-fold assessments are based on the samples in the left-out fold at each step of the cross-validated analysis. The training set yielded an out-of-fold performance metric, Area Under Curve (AUC), of 0.96 (Elastic net and Lasso) with an internal sample test AUC of 0.84 (Elastic net) and 0.88 (Lasso) (Figure 4A). The distribution of probability scores indicated that within the training data both models classify well (4B) but that improved robustness and stability of scoring was found with the Elastic net model as evidenced by reduced variation in probability scores observed during repeated cross-validations. Next, the training model was tested on two external validation set of patient samples. These include pancreatic cancer and non-cancer samples from Li et al^24^ (pancreas subtype was not specified, 23 pancreas, 53 non-cancer) and Song et al^22^ (pancreas subtype specified as adenocarcinoma, 7 pancreas, 10 non-cancer – Supplementary Figure 2). The validation set exhibited a performance with AUC of 0.78 (Elastic net and Lasso) in the Li et al data and AUC of 0.99 (Elastic net) and 0.97 (Lasso) in the Song et al data (Figure 4C).

**Figure 4:**
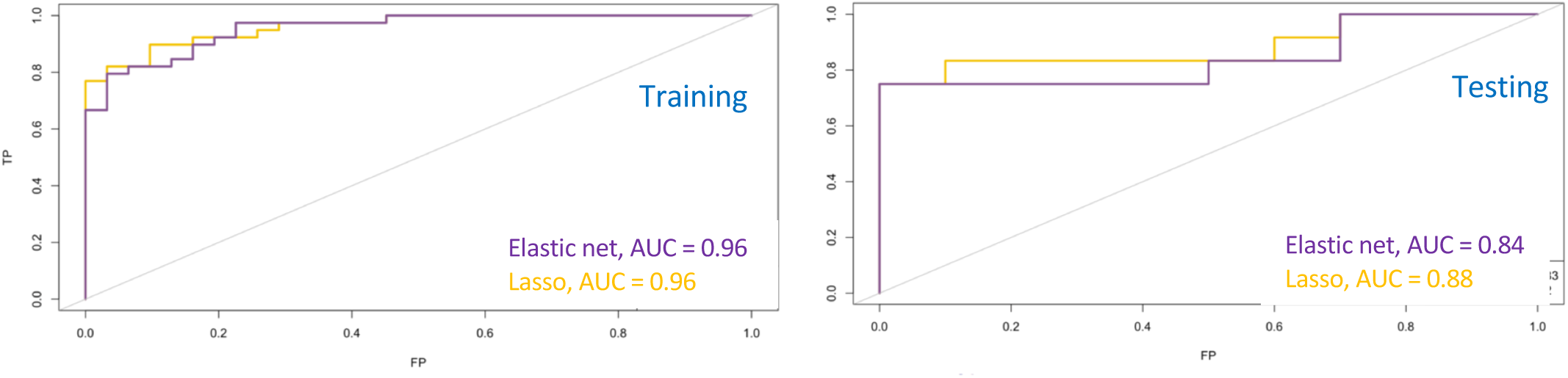
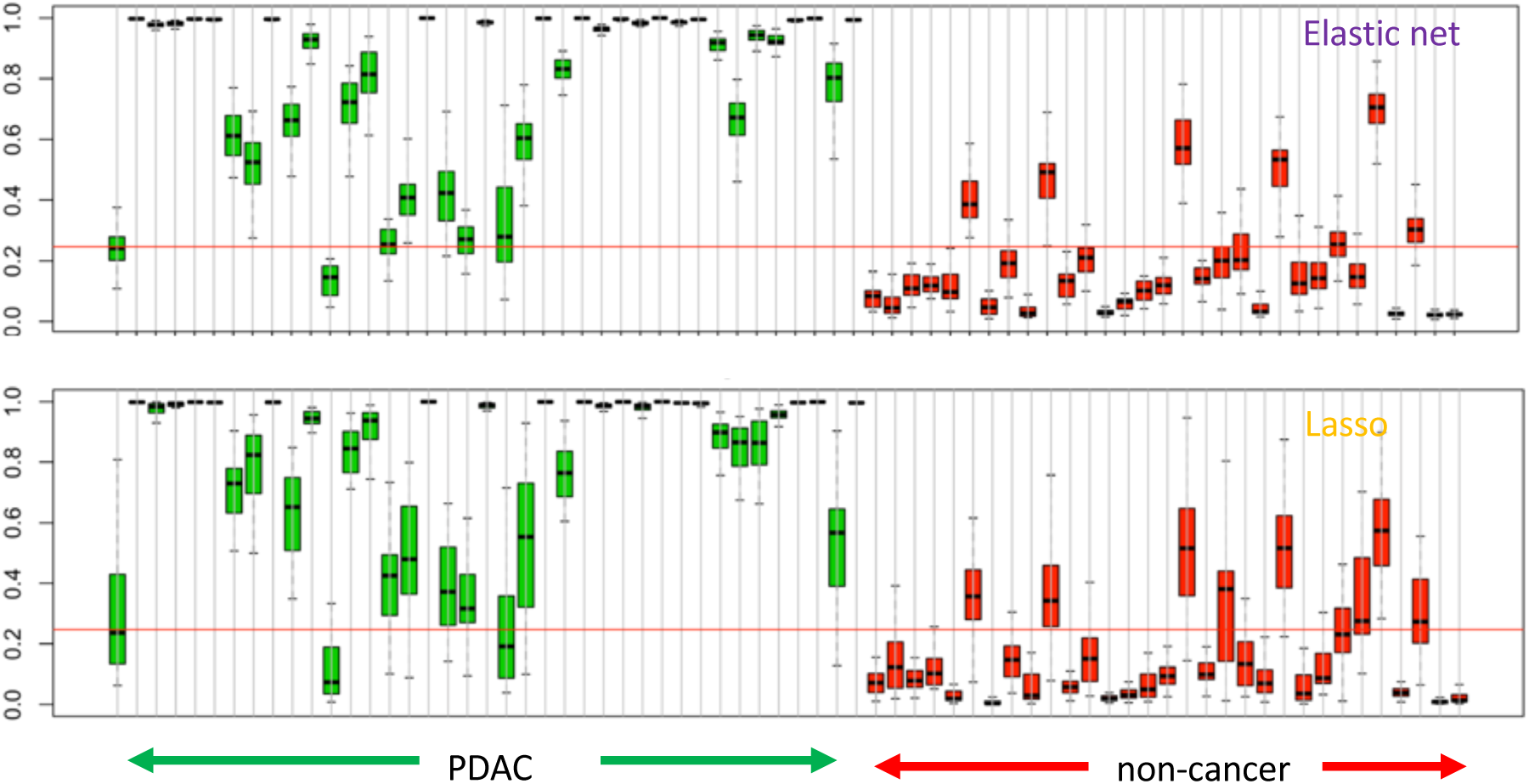
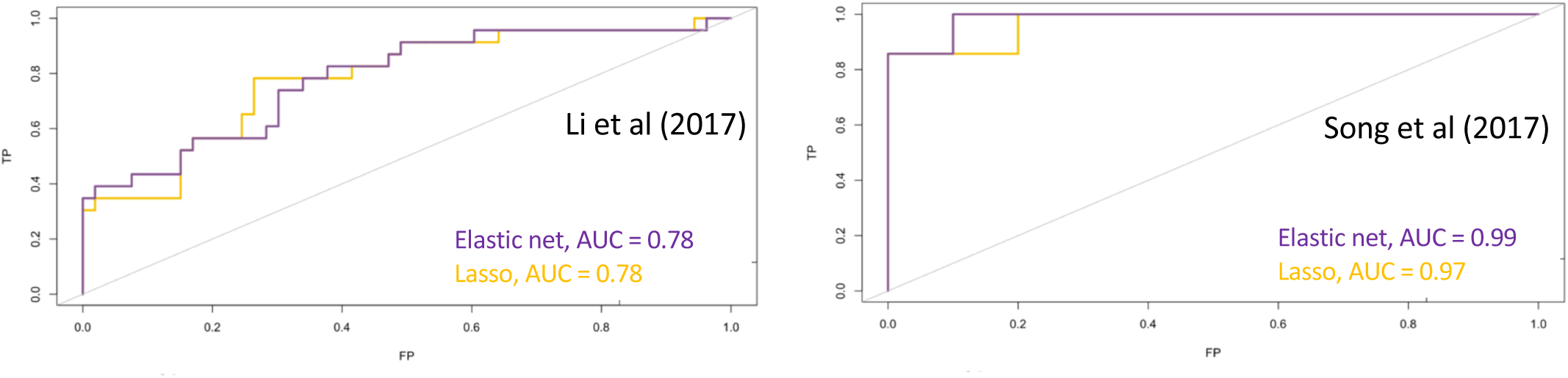
Identification of a 5hmC signature that differentiates PDAC from non-cancer samples. A. Predictive modeling using two regularization models (Elastic net and Lasso) on 75% of the data (training) data - left panel. Test performed on the remaining 25% of original data – right panel. B. Probability scores derived for each sample in the training dataset using the Elastic net and Lasso regularization models. Probability scores towards 1 are predicted cancer samples whereas probability scores close to 0 are non-cancer samples. Red line – identified Q3 probability score of the non-cancer samples. C. Validation of predictive models using Li et al (2017) and Song et al (2017) pancreas and healthy sample data sets.

The effect of feature selection on prediction performance was evaluated by filtering the initial set of significant genes (Figure 3A) to satisfy a 1.5 fold change in 5hmC representation in the PDAC cohort with median gene count of log2 average > 4. This filtering approach was applied on 75% of sample data, reserving the remaining 25% for subsequent testing (see below). The same regularized regression models were built using the resulting set of 287 and 343 genes with increased and decreased 5hmC density, respectively. Employing a similar setup for training and testing as defined previously, we found the training set produced an AUC of 0.96 (Elastic net) and 0.94 (Lasso). Not surprisingly, internal testing yielded a high performance with AUC of 0.92 (Elastic net) and 0.93 (Lasso). Of greater interest, was the performance on external data sets with AUC of 0.74 (Elastic net) and 0.67 (Lasso) for Li et al data and AUC of 0.97 (Elastic net) and 0.94 (Lasso) for Song et al data. This suggests that statistically filtered genes that are biologically relevant to pancreatic cancer and/or pancreas development do not perform much better than an algorithmically driven selection of features during regression training, as has been shown elsewhere^32^.

The hyper-parameters were discovered from the models built by the analysis of the 65% most variable 5hmC gene features in the training set. These hyper-parameters were employed to fit the full cohorts of PDAC and non-cancer samples and yielded models with 109 genes (Elastic net) and 47 genes (Lasso). Genes used in the models were found to possess PDAC versus non-cancer t-scores that are concordant with both Li et al and Song et al data sets (Figure 5).

**Figure 5.**
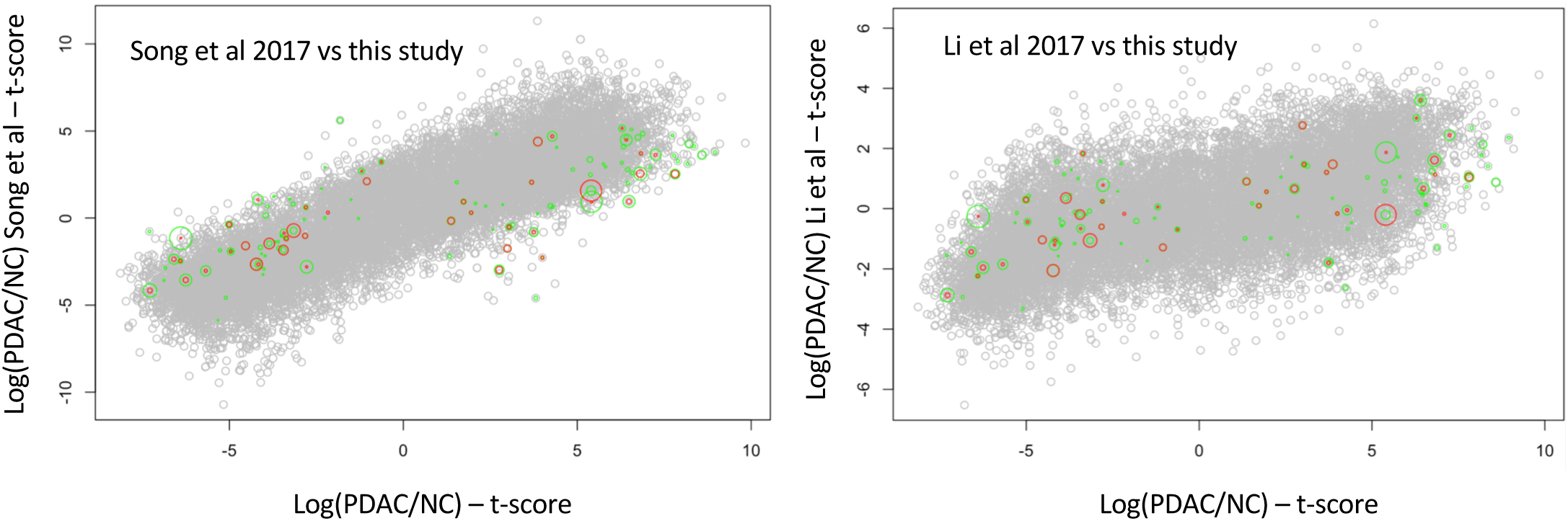
Comparison of t-scores of 5hmC density fold difference between PDAC and noncancer (NC) cohorts as found in (A) Song et al and (B) Li et al, each compared to this study. All gene scores are represented in grey, Elastic net model genes are in green and Lasso model genes are in red. The size of each green and red dot represents the relative contribution of that gene to the model.

## Discussion

This study was focused on the discovery of cfDNA specific hydroxymethylation-based biomarkers that may facilitate the development of molecular diagnostic tests to detect pancreatic cancer at earlier stages. Our data highlight the ability to detect differentially hydroxymethylated genes whose underlying biology shows association with both pancreas and cancer development as well as established trends in chromatin mark maps and other functional regions of the genome. Furthermore, regularized regression methods were used to build models from (i) statistically filtered genes that form a biologically relevant set and (ii) a comprehensive gene set that is found to be highly variable, yielding models with AUC of 0.94-0.96 along with an external data set validation AUC of 0.74-0.97 (Elastic net models). The 5hmC signal was readily found to overlap in gene-centric functional regions (enrichment in promoter, exons, UTR and TTS), as well as transposable elements like SINEs (enriched) and LINEs (depleted) (Figure 2A, B). Such hydroxymethylcytosine changes in functional regions have been reported in cfDNA from colorectal^24^, esophageal^23,33^ and lung cancer^23^. In a similar manner, PDAC specific gains or losses in hydroxymethylation were observed in functional regions in our data. In addition to enrichment and depletion of 5hmC in functional regions, there was a novel PDAC specific 5hmC increase in exons, TTS and 3’UTR and 5hmC decrease in promoters and LINE elements (Figure 2 C). In embryonic stem cells, 5-hydromethylation decreases in the promoter region have been shown to associate with elevated gene transcription^34^. An increase in disease relevant transcription may be implicitly supported in our PDAC data by the 5hmC increase in gene-centric features mentioned earlier, as well as an apparent decreasing trend of 5hmC in promoter regions toward late stage PDAC (Figure 2 D, E). Dynamic changes in chromatin have been shown to control cell development and transition of cells with oncogenic potential^35^. Intersection of our 5hmC data with histone maps of the pancreatic cancer cell line PANC-1 available from ENCODE project revealed 5hmC decrease in PDAC cfDNA (Figure 2 E). Consistent with 5hmC changes over promoters, 5hmC decrease over H3K4me3 loci also suggests disease specific increases in gene transcription via chromatin modifications, given the permissive transcriptional state associated with H3K4me3^36^. 5hmC patterning around known functional elements of the genome suggests a broader interplay between hydroxymethylation and the epigenetic control of transcriptional processes. Additional work will reveal the extent to which models predictive of PDAC can be built from a combination of gene-specific features, genomic loci with different chromatin states and transposable elements detected in cfDNA.

In this study, we employed coarse resolution of hydroxymethylation at gene-based level in PDAC, and yet, were able to find genes whose increased 5hmC signals highlighted pathways implicated in liver cancer (2). While it remains to be investigated whether the observation of pathways implicated in liver cancer is due to PDAC burden on liver, we note that pancreas typically has groups of expressed genes shared with liver and salivary gland (https://www.proteinatlas.org/humanproteome/pancreas) and MSigDB does not currently contain pathways annotated for pancreatic cancer^37^.

To further investigate biological functions possibly regulated by 5-hydroxymethylation changes identified herein for PDAC, we employed two approaches for gene set enrichment analysis; (i) using genes with differentially decreased 5hmC or (ii) via performing GSEA on all reporting genes., which implicated close to one third of immune system pathways. Assuming a positive correlation between 5hmC density in gene bodies and gene transcription, as was previously reported, one interpretation of this result is that immune system function is decreased in PDAC patients.

Inspection of individual genes that were either significantly increased or decreased in 5hmC density revealed genes implicated in normal pancreas development, for instance the transcription factors GATA4, GATA6, PROX1, ONECUT1/2, in addition to genes whose increased expression is implicated in cancer, such as YAP1, TEAD, PROX1, ONECUT2, ONECUT1, IGF1 and IGF2. The relative 5hmC increase in transcription factor genes like GATA4, GATA6, PROX1, ONECUT1/2, which were previously reported to be involved in early pancreatic development^26–28^, suggest a reversion to a stem-like state in PDAC samples. This is further supported by the identification of a stem cell pathway (BOQUEST_STEM_CELL_UP) among the top 20 mSigDB pathways determined using 142 genes with increased 5hmC in PDAC cfDNA.

Identifying genes whose 5hmC densities are significantly changed in PDAC, leads to an enrichment of genes with annotated relevant biology which can be used to build regularized regression models, whose performance matched models built on the more comprehensive set of variable genes. This gives us good confidence that our models, whose performance is high (training AUC of 0.94-0.96 with an external data set validation AUC of 0.74-0.97), are measuring underlying biological signals relevant to PDAC. We note that our current external data set performance may also be impacted by the small sample size of external data sets (Li et al, 23 pancreas, 53 healthy and Song et al, 7 pancreas, 10 healthy). Moreover, whilst our external validation AUC on Li et al data was generally lower than the one for Song et al data, we note that differences in PDAC sample distribution over various cancer stages may contribute to this finding. While Song et al was broadly distributed across all stages, Li et al pancreatic data were distributed with a mode around stage 3 disease versus a mode at stage 2 for this study (Supplementary Figure 2). Therefore, our predictive model may be better suited for the detection of earlier versus later stage disease. Other patient characteristics (such as histological subtype and smoking, etc.) may have also differed in these independent study sets.

Despite the large number of differentially hydroxymethylated genes, the regularized regression models included 100 genes or less. However, the fact that 13,180 differentially hydroxymethylated genes were detected in PDAC cfDNA compared to non-cancer cfDNA suggests that other biological signals may also reside in our data set. Smoking status is a known risk factor for PDAC up to 20 years post smoking cessation and DNA methylation changes have been associated with tobacco-based toxins^38^. In our retrospective case-control designed study, ever smokers constituted 59% and 49% of PDAC and non-cancer cohorts respectively, indicating that ever smokers are well represented in each cohort. Consequently, we do not suspect that smoking association in our PDAC cohort could account for the significantly hydroxymethylated genes found. However, a more extensive future study focused on sub-partitioning PDAC and non-cancer patient into never and ever smokers with pack-year characteristics will enable us to address the impact of smoking on the hydroxymethylome in PDAC patients. These pancreatic cancer risk parameters combined into a clinically relevant, intent-to-test population-based study, will further our findings beyond our current case-control cohort study, which numbers less than 100 participants. Further consideration of disease-related clinical parameters will enable us to explore hydroxymethylcytosine features with the aim of yielding refined signals capable of earlier diagnosis of PDAC.

## Methods

### Clinical cohorts and study design

– A case-control study was performed using plasma obtained from subjects without (termed non-cancer) and with pancreatic cancer who provided informed consent and contributed biospecimens in studies approved by the Institutional Review Boards (IRBs) at participating sites in the United States and Germany. Plasma samples for the non-cancer cohort were obtained from subjects enrolled prospectively at five sites in the United States, following review and approval of the study protocol by each site’s participating investigator(s).

### Cancer cohort

- Plasma samples for the cancer cohort were obtained from subjects who had undergone management for pancreatic cancer in the United States or Germany, and also provided consent for use of blood specimens for archival storage and retrospective analyses. Criteria for subject eligibility for inclusion in the analysis included age greater than or equal to 21 years for all subjects, with additional requirements for the cancer cohort including: 1) no cancer treatment, e.g., surgical, chemotherapy, immunotherapy, targeted therapy, or radiation therapy, prior to study enrollment and blood specimen acquisition; and 2) a confirmed pathologic diagnosis of adenocarcinoma inclusive of all subtypes.

### Non-cancer cohort

- Subject exclusion criteria for the non-cancer cohort also included any of the following: prior cancer diagnosis within prior six months; surgery or invasive procedure requiring general anesthesia within prior month; non-cancer systemic therapy associated with molecularly targeted immune modulation; concurrent or prior pregnancy within previous 12 months; history of organ tissue transplantation; history of blood product transfusion within one month; and major trauma within six months. Clinical data required for all subjects included age, gender, smoking history, and both tissue pathology and grade, and were managed in accordance with the guidance established by the Health Insurance Portability and Accountability Act (HIPAA) of 1996 to ensure subject privacy.

### Plasma collection

- Plasma was isolated from whole blood specimens obtained by routine venous phlebotomy at the time of subject enrollment. For cancer subjects, whole blood was collected in K3EDTA tubes (Sarstedt, Nümbrecht, Germany) with isolation of plasma within 4 h of phlebotomy by centrifugation at 1,500g for 10 min at RT, followed by transfer of the plasma layer to a new tube for centrifugation at 3,000g for 10 min at RT, with plasma aliquots used for isolation of cell-free DNA (cfDNA) or stored at −80°C.

For non-cancer subjects, whole blood was collected in Cell-Free DNA BCT^®^ tubes according to the manufacturer’s protocol (Streck, La Vista, NE) (https://www.streck.com/collection/cell-free-dna-bct/). Tubes were maintained at 15 °C to 25 °C with plasma separation performed within 24 h of phlebotomy by centrifugation of whole blood at 1600 × g for 10 min at RT, followed by transfer of the plasma layer to a new tube for centrifugation at 16,000 × g for 10 min. Plasma was aliquoted for subsequent cfDNA isolation or storage at −80°C.

### cfDNA isolation

– cfDNA was isolated using the QIAamp Circulating Nucleic Acid Kit (QIAGEN, Germantown, MD) following the manufacturer’s protocol excepting the omission of carrier RNA during cfDNA extraction. Two milliliter plasma volumes (cancer) or four milliliter plasma volumes (non-cancer) were lysed for 30 minutes prior to collection of nucleic acids; all cfDNA eluates were collected in a volume of 60 μl buffer. All cfDNA eluates were quantified by Bioanalyzer dsDNA High Sensitivity assay (Agilent Technologies Inc, Santa Clara, CA) and Qubit dsDNA High Sensitivity Assay (Thermo Fisher Scientific, Waltham, MA) was employed to ensure the absence of contaminating high molecular weight DNA emanating from white blood cell lysis.

### 5-hydroxymethyl Cytosine (5hmC) assay enrichment

– Sequencing library preparation and 5hmC enrichment was performed as described previously (Song et al). cfDNA was normalized to 10 ng total input for each assay and ligated to sequencing adapters. 5hmC bases were biotinylated via a two-step chemistry and subsequently enriched by binding to Dynabeads M270 Streptavidin (Thermo Fisher Scientific, Waltham, MA). All libraries were quantified by Bioanalyzer dsDNA High Sensitivity assay (Agilent Technologies Inc, Santa Clara, CA) and Qubit dsDNA High Sensitivity Assay (Thermo Fisher Scientific, Waltham, MA) and normalized in preparation for sequencing.

### DNA sequencing and alignment

– DNA sequencing was performed according to manufacturer’s recommendations with 75 base-pair, paired-end sequencing using a NextSeq550 instrument with version 2 reagent chemistry (Illumina, San Diego, CA). Twenty four libraries were sequenced per flowcell and raw data processing and demultiplexing was performed using the Illumina BaseSpace Sequence Hub to generate sample-specific FASTQ output. Sequencing reads were aligned to the hg19 reference genome using BWA-MEM with default parameters^39^.

## Peak Detection

BWA-MEM read alignments were employed to identified regions or peaks of dense read accumulation that mark the location of a hydroxymethylated cytosine residue in a CpG content. Prior to identified peaks BAM files containing the locations of aligned reads were filtered for poorly mapped (MAPQ < 30) and not properly paired reads. 5hmC peak calling was carried out using MACS2 (https://github.com/taoliu/MACS) with a p-value cut off = 1.00e-5. Identified 5hmC peaks residing in “blacklist regions” as defined elsewhere (https://sites.google.com/site/anshulkundaie/proiects/blacklists) and read date on chromosomes X, Y and mitochondrial genome were also removed. Computation of genomic feature enrichment overlap 5hmC peaks were performed using the HOMER software (http://homer.ucsd.edu/homer/) with default parameters.

Chromatin modifications (H3K4me1, H3K4me3 and H3K27ac) were identified in histone maps of the pancreatic cancer cell line PANC-1 and were downloaded from ENCODE ChIP-Seq repository (https://genome.ucsc.edu/encode/dataMatrix/encodeChipMatrixHuman.html Determination of enrichment were calculated via Odds ratio using the Fisher Exact Test via the program bedtools fisher. For comparisons between PDAC and non-cancer, the Wilcoxon test was used, and for across stages comparison, the Kruskal-Wallis Test was employed.

## Differential Representation Analysis

For the purpose of reliably identifying gene bodies with differential representation between the PDAC and the non-cancer groups, we closely followed the RNA-Seq workflow outlined in Law et al. 2016^40^, including much of the preliminary QC steps. In brief, the analysis includes data pre-processing by adopting the following workflow: (i) transforming the data from raw counts to log2(counts per million), (ii) removing genes that are weakly represented, (iii) normalizing the gene representation distributions, and (iv) performing unsupervised clustering of samples. To accomplish differential representation analysis, we applied the following steps: (i) creating a design matrix and PDAC vs non-cancer cohorts, (ii) removing heteroscedascity from the data, (iii) fitting the linear models for the comparison of interest, PDAC vs non-cancer, (iv) examining the number of differentially represented genes.

In most of these analysis steps the default settings were used when appropriate. To remove weakly represented genes, we excluded genes that did not have greater than 3 counts per million reads in at least 20 samples. This filter excludes roughly 12% of the genes. For the identification of the significantly differentially represented regions we used the method of Benjamini and Hochberg^41^ to obtain p-values adjusted for multiple comparisons. In this report we use adjusted p-value and false discovery rate (FDR) interchangeably.

## Predictive Modelling

For the purpose of assessing the feasibility of building classifiers that can discriminate between PDAC and non-cancer samples based on the 5hmC representation of gene bodies, we evaluated the performance of two forms of regularized logistic regression models, namely the Lasso and the Elastic net, which are commonly used in the classification context, where the number of examples are few and the number of features are large. See Friedman et al. (2010)^31^ for a description of the general Elastic net procedure. Software implementation of these methods can be found at https://cran.r-project.org/web/packages/glmnet/index.html. Weakly represented genes were excluded from analysis as described in the section on **Differential Representation Analysis**.

All training and fitting were done on 75% of the samples selected at random in a balanced way to keep the ratio of the number of PDAC to non-cancer samples similar in both the training and testing subsets. Before any fitting, genes were filtered to include the 65% of the most variable genes for model fitting task. The filter was designed using training samples only and was done in a way to ensure that genes of all levels of 5hmC representation were included.

Both assessed regularization methods, the lasso and the elastic net, require specifying hyper-parameters which control the level of regularization used in the fit. Hyper-parameters were selected based on out-of-fold performance on 30 repetitions of 10-fold cross-validated analysis of the training data. Out-of-fold assessments are based on the samples in the left-out fold at each step of the cross-validated analysis. The out-of-fold performance of the models fitted with hyper-parameter values set at the optimal values might yield a slightly optimistic assessment of performance. The performance of these models applied to the test set should provide less biased estimate of performance, although generalizability to external datasets is not always guaranteed.

The hyper-parameter values that lead to the best out-of-fold performance were then used to fit the final models which were fitted to the entire set of samples including both training and testing subsets. The performance of these final models can thus only be evaluated based on their performance on external data sets. These do provide a sense of the generalizability of the performance observed in the local training and testing data sets.

To evaluate the effect of feature selection on prediction performance, we repeated the training and evaluation task based on a filtered set of genes that included genes found to be significantly differentially represented, having a 1.5 fold differential 5hmC representation, and a level of representation exceeding the median level (log2 CPM ≥ 4). This filter was designed based on training data statistics only.

---

**Supplementary Figure 1**. Jaccard index population frequency plots showing similarity of H3K4me1, H3K4me3 or H3K27ac histone maps with 5hmC peaks in Non-Cancer (NC) and Pancreas (PDAC) cohorts. Frequency in y-axis calculated as: density x number of points/sum of density x number of points.

**Supplementary Figure 2**. Distribution of patients over pancreatic cancer stages I-IV from each study (BSG: Bluestar genomics cohort, Li et al (2017) and Song et al (2017)) expressed as fraction of cohort.

